# A screening platform based on epitope editing for drug discovery

**DOI:** 10.1101/838896

**Authors:** Biyue Zhu, Jing Yang, Richard Van, Kathleen Ran, Keyi Yin, Yingxia Liang, Xunuo Shen, Wei Yin, Se Hoon Choi, Ying Lu, Changning Wang, Yihan Shao, Rudolph E. Tanzi, Can Zhang, Yan Cheng, Zhirong Zhang, Chongzhao Ran

## Abstract

The interaction between an antibody and its epitope has been daily utilized in various biological studies; however it has been rarely explored whether small molecules can alter the interaction. We discovered that small molecules could alter/edit surface properties of amyloid beta (Aβ) epitopes, and consequently inhibit or enhance corresponding antibody recognition. Remarkably, this editing effect could generate functional changes including protein aggregation behaviors, cell cytokine secreting and in vivo microglia activation. According to this discovery, we proposed a <u>s</u>creen <u>p</u>latform based on <u>e</u>pitope <u>e</u>diting for <u>d</u>rug discovery (SPEED). With a small library of compounds, we validated that SPEED could be used to seek new leads for Aβ species. We also demonstrated that this platform could potentially be extended to other targets including tau protein and PD-L1 protein. The SPEED is a simple, fast and label-free screening method. We believe that the SPEED strategy could be universally applicable for seeking and validating drug candidates and imaging ligands.

## Introduction

Genes and proteins are the most essential components linked with functions of a living organism. While genes encode information, proteins execute biological functions. Genome editing, which enables adding, removing, or altering nucleotide sequences at specified positions, is currently considered as one of the most powerful tools for fundamental biological studies and clinical research.^1, 2^ In parallel with genome editing, the alteration of amino acid sequence and hierarchical structures could be considered as editing action of a protein. Numerous approaches to modify/edit the amino acid sequence have been reported and explored for studying basic mechanism and for seeking therapeutics.^3, 4^ For the hierarchical structures of a protein, surface properties including charge/electrostatic force and hydrophobicity/Van der Waals forces are the primary determinants. Alternation of charge and hydrophobicity of an amino acid sequence of interest could also be deemed an editing process. For an antigen of a protein/peptide, an epitope is normally the antibody binding sequence of amino acids, and it can be the whole sequence or a fragment.^5^ Although the epitope-antibody interactions have been extensively applied in various biological studies, it has been rarely explored whether the interaction can be altered by inserting an additional molecule. For the binding between an antigen with its antibody, charges and hydrophobicity are the major interacting forces.^6^ We speculated that, when a molecule (editor) bind to an epitope, the readout of corresponding antibody binding could be different due to the altering/editing of the surface properties. We posited that this phenomenon could be used as a screening strategy to determine whether the molecule (editor) can bind to the epitope (protein) and reflecting the degree of epitope-drug interactions. We termed it “<u>s</u>creening <u>p</u>latform based on <u>e</u>pitope <u>e</u>diting for <u>d</u>rug discovery (SPEED)”. To the best of our knowledge, such investigation has been so far neglected.

In the present study, we demonstrated that SPEED could be used to establish a versatile screening platform to seek and validate ligands, therapeutics, and imaging probes. Amyloid beta (Aβ), one of the major hypothesized pathogenic targets in Alzheimer’s disease (AD), was first selected as the model protein for proof-of-concept studies, in which we investigated the altering/editing effect of charge and hydrophobicity of Aβ fragment on antibody binding. Next, we investigated functional changes after altering/editing Aβ epitope, include protein aggregation behaviors, cytokine secretion from microglial cells and in vivo microglial activation. With very encouraging results, we explored whether SPEED could be used to screen for Aβ leads with a small library of 32 compounds. Lastly, we used tau protein and PD-L1 protein as two additional examples to further demonstrate the potential applicability of SPEED.

The SPEED strategy requires no modification or tagging of the tested protein and the drug candidate, and can be easily and widely implanted into many biomedical laboratories without extra equipment. We believe that SPEED is an important complement for the currently available drug screening approaches.

## Results

### Experimental design for proof-of-concept

Over the past decade, our research group has been working on imaging probe development for Aβ species. In the course of our studies, we observed that some molecules could potentially alter the surface properties of certain fragments of Aβ peptides. Based on the observations, Aβ was selected as the model protein for proof-of-concept studies. Additional considerations include: 1) Aβ is one of the primary hypothesized pathogenic targets in AD and was reported to correlate with neuroinflammation and production of auto-antibodies.^7^ Clearly Aβs could be considered as an antigen and its fragment as an epitope; 2) Surface properties of Aβ play a major role in protein assembly and are closely associated with its toxicity, enzyme activation, oxidative stress and other functions;^8, 9^ and 3) Aβ already has commercially available antibodies for various known epitopes.^10^

Next, we set out to choose a proper method for target protein immobilization. Rather than using resins, capture antibodies, and functional groups to trap/react with target proteins, we used dot blot immunoassay as the screening platform for straightforward investigation due to: 1) no extra modification of protein is required, which could maintain the original structure properties of proteins; 2) the changes of target epitope can be easily monitored by amplified fluorescence and chemiluminescence signals from antibodies (the most widely used technique for readout of Western blot and ELISA); 3) the methodology validations of dot blot have already been well-established, showing high sensitivity with detecting limit as low as picogram quantities. With rapid and easy procedures, dot blot enables dozens of assays on one assay strip; 4) the inexpensive apparatus enable fast immobilization of proteins that could be easily and widely implanted into many biomedical laboratories.^11, 12^ In our studies, the screening platform was successfully set up via the following procedure: protein/peptidic antigen with or without drug/editor treatment was immobilized on a nitrocellulous membrane using Bio-Dot Microfiltration Apparatus. Each strip including 5-6 duplications was assigned to corresponding antibody recognition for epitope of interest (EOI). The chemiluminescence readout from control group (without tested drug/editor) was normalized to 1.0, and the editing effect for EOI was quantified using the ratio between the two groups.

### Charge/Electrostatic Editing of Aβs

Charge/electrostatic interactions on an epitope play critical roles for antibody recognition by cation–π interaction, anion–π interactions and hydrogen bond formation.^6^ To investigate whether modulating electrostatic interactions on an epitope could lead to changes of antibody-binding readout, we choose 12-crown-4 ether as a model compound for charge/electrostatic editing. Previously we found that 12-crown-4 could attenuate the aggregation of Aβs.^13^ Reportedly, crown ethers could form hydrogen bonds with positively charged amino groups (e.g. lysine, arginine, histidine) to shield the charge,^14^ and the sequence of Aβ peptides contains important lysine (K16), a crucial amino acid that forms inter-sheet salt bridges and promotes mis-folding of Aβ.^9^ Thus, we hypothesized that 12-crown-4 could interact with the positively charged e-amino of K16 via forming hydrogen bonds and this interaction could alter/edit the surface charge of Aβ, which could consequently interfere the binding between epitope and corresponding antibody (Fig. 1a).

**Fig.1.**
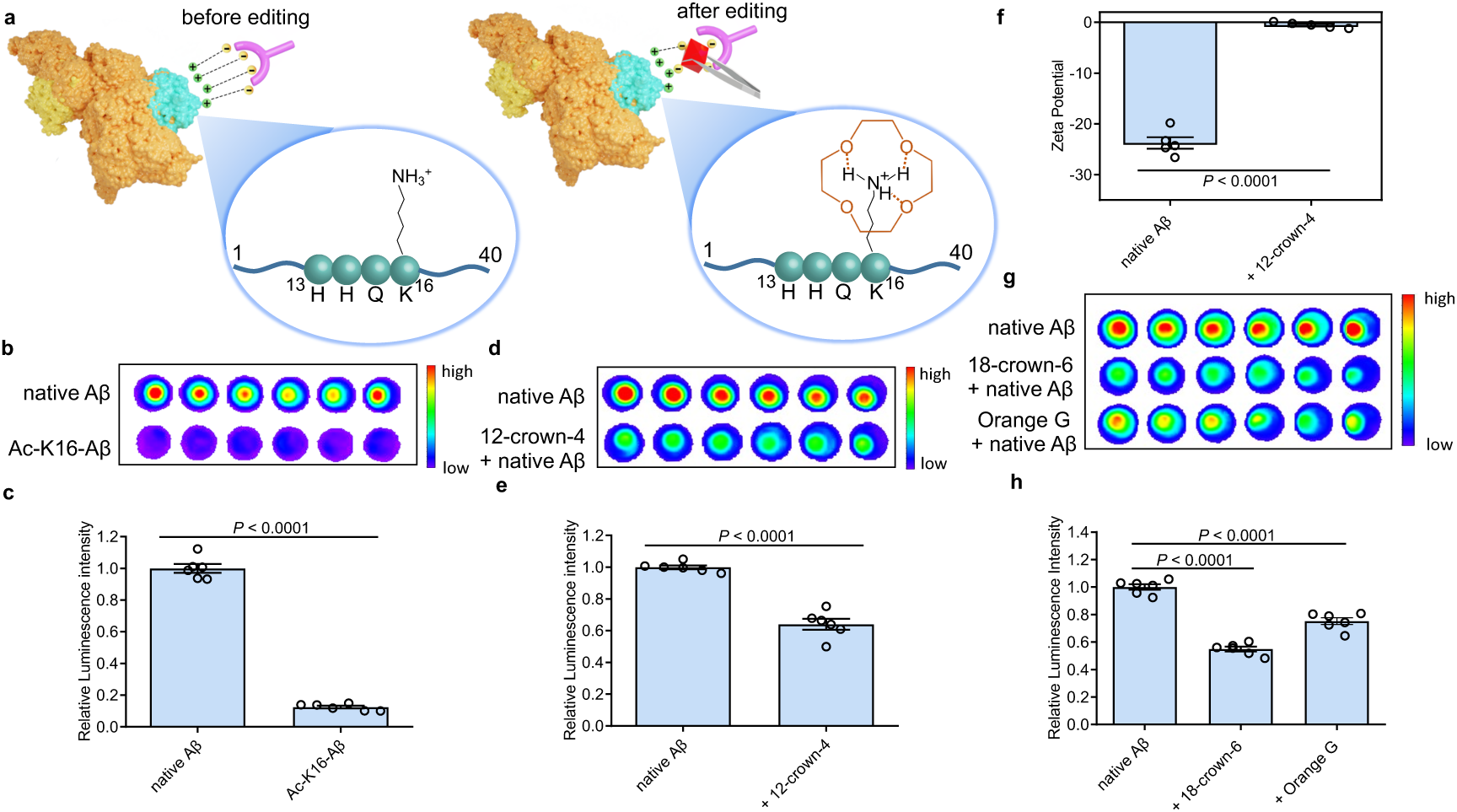
Charge/Electrostatic editing effect. **a)** Illustration of the interaction between an epitope and its corresponding antibody before and after charge/electrostatic editing. **b,c**) SPEED assay (**b**) and quantitative analysis (**c**) for native Aβs and acetylated-K16 Aβs (Ac-K16-Aβ) using 6E10 antibody (n = 6). Ac-K16-Aβs showed decreased signal compared with non-modified Aβs, indicating the positive charge on K16 is critical for 6E10 antibody recognition. **d,e**) SPEED assay (**d**) and quantitative analysis (**e**) for native Aβs with or without 12-crown-4 using 6E10 antibody (n = 6). The addition of 12-crown-4 showed decreased signals of native Aβs compared to the non-treated group, suggesting 12-crown-4 was capable of altering the charge status of the epitope. **f**) Zeta potential of Aβs with or without 12-crown-4. The zeta potential of Aβs significantly increased after interacting with 12-crown-4, further confirming the charge editing of the epitope. **g,h**) SPEED assay and quantitative analysis for native Aβs with or without 18-crown-6 and Orange G using 6E10 antibody (n = 6). 18-crown-6 and Orange G showed similar editing effect on binding as 12-crown-4. Data in **c,e,f,h** are mean ± s.e.m. and *P* values were determined by two-tailed Student’s *t*-test.

First, to validate that modification of surface charge of the Aβ peptide can alter antibody binding, e-amino of K16 was acetylated (Ac-K16-Aβ) to turn positive K16 into neutral, and the chemiluminescence readouts from native Aβ peptide and Ac-K16-Aβ were compared using 6E10 antibody, which is reactive to 1-16 amino acids (a.a.). Indeed, much lower intensity was observed from Ac-K16-Aβ group than that from the non-modified Aβ group (Fig.1b,c), indicating the positive charge on K16 is critical for the binding between Aβ and 6E10 antibody. Next, 12-crown-4 was used to investigate whether it could have similar effects on the binding as the acetylation of K16. After mixing 12-crown-4 with peptides (native Aβ and Ac-K16 Aβ), we found that the intensity of native Aβ was significantly decreased, while no obvious signal change of Ac-K16 Aβ was observed (Fig.1d,e and Supplementary Fig.1a), suggesting that 12-crown-4 was capable of altering the charge status of the epitope. The altering of surface charge of Aβ by 12-crown-4 was further validated by zeta potential assays, which have been regularly used to measure changes of surface charge status.^15^ As expected, the zeta potential of Aβ significantly increased after interacting with 12-crown-4, further confirming the charge/electrostatic editing within the epitopes (Fig.1f).

Based on the encouraging results from the 12-crown-4 study, we set out to test its analogues Icotinib and 18-crown-6. 18-crown-6 showed similar editing effect on binding as 12-crown-4 while Icotinib showed no significant editing effect (Fig.1g,h and Supplementary Fig.1b). This may be due to the structure of Icotinib, which is relatively bulky to interact with the core of Aβs. Interestingly, 12-crown-4 and 18-crown-6 could only reduce the readout of aggregated Aβs but not monomeric and oligomeric Aβs (Supplementary Fig.1c). This is likely due to the positive charges buried inside the Aβ monomers and oligomers, which have less ordered transient states.^16^ Next, we used Orange G as an additional example for editing K16, as previous X-ray analysis of atomic structure indicated the negatively charged sulfonic acid groups of Orange G could interact with positively charged K16.^9^ As expected, Orange G showed similar results as crown ethers (Fig.1g,h). Collectively, our results indicated that the positive charge on K16 was essential for the antibody binding and it could be altered/edited by crown ethers and Orange G.

### Hydrophobicity Editing of Aβs

It is estimated that paratopes, the antigen-binding site on antibodies, are enriched with aromatic residues (Tyr, Trp, and Phe), especially those with short hydrophilic side chains (Ser, Thr, Asp, and Asn).^6^ Previously we reported that curcumin analogue CRANAD-17 could bind to Aβ species and attenuate cross-linking of Aβs, and ^1^H NMR spectra showed that CRANAD-17 could interact with Aβ16-20 fragment, which is the core fragment of Aβ peptides.^17^ The curcumin scaffold of CRANAD-17 serves as the anchor moiety for targeting the hydrophobic core of Aβ, and we hypothesized that the benzene and imidazole rings on both sides of CRANAD-17 could altered/edited hydrophobic properties around Aβ17-24 epitope, which can be recognized by 4G8 antibody (Fig. 2a). To this end, the chemiluminescence readouts from Aβs with or without CRANAD-17 were compared using 4G8 antibody. Thioflavin T (ThT), a standard dye for Aβ plaques but not specific to Aβ17-24,^18, 19^ was used as a negative control. We found that CRANAD-17 group displayed significantly enhanced chemiluminescence intensities for all Aβ species (Fig.2b,c and Supplementary Fig.2 for oligomers and aggregates). Although we expected CRANAD-17 to change the readouts, the increase in chemiluminescence intensity was surprising. For ThT, our data showed no apparent changes of Aβ species with and without ThT, which is in accordance with previous study that ThT primarily binds in channels running parallel to the long axis of Aβ fibrils (Supplementary Fig.3).^18^ To investigate the specificity and sensitivity of this method, we used Aβ 6E10 antibody (reactive to Aβ1-16) for comparison. As shown in Fig.2b-e, CRANAD-17 displayed more robust increase in signal from 4G8 antibody than that from 6E10 antibody, confirming that the editing effect of CRANAD-17 is specific on epitope Aβ17-24. Furthermore, we performed a concentration-dependent titration and using 4G8 antibody for detection. As shown in Fig.2f and Supplementary Fig.4, the enhancement was in a concentration-dependent manner, and the EC50 was about 121.0 nM.

**Fig.2.**
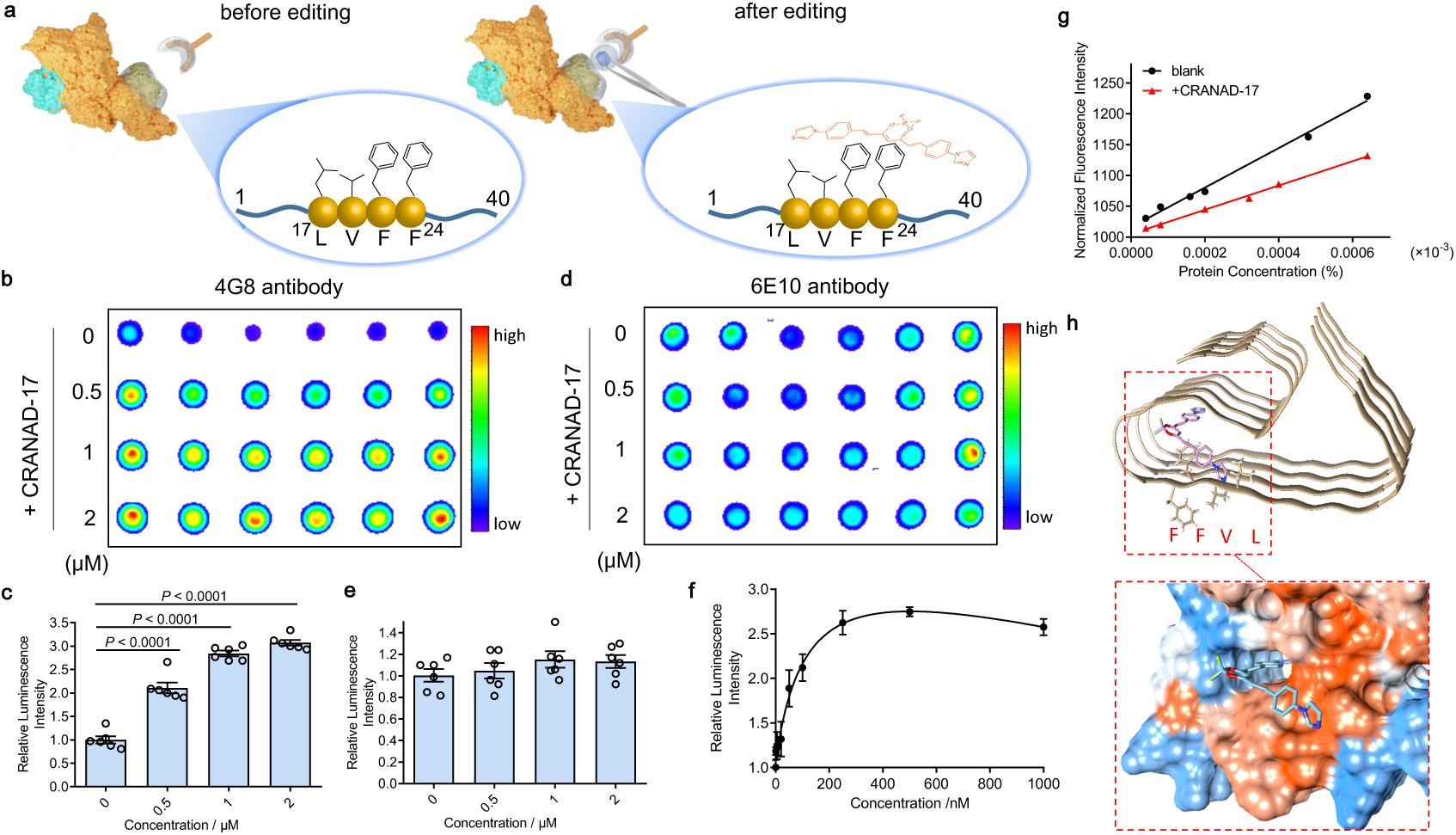
Hydrophobicity editing effect of CRANAD-17. **a)** Illustration of the interaction between an epitope and its corresponding antibody before and after hydrophobicity editing. **b-e**) SPEED assays and quantitative analysis for Aβs with different concentrations of CRANAD-17 by using 4G8 (**b,c**) and 6E10 (**d,e**) antibody (n = 6). CRANAD-17 displayed significantly enhanced chemiluminescence signals from 4G8 antibody, which is more robust than that from 6E10 antibody, indicating the editing effect of CRANAD-17 is specific on epitope Aβ17-24. Data in **c,e** are mean ± s.e.m. and *P* values were determined by two-tailed Student’s *t*-test. **f**) Concentration dependent profile of CRANAD-17 (1-1000 nM) with Aβs (1 μM) determined by SPEED assay. Half maximal effective concentration (EC50 = 121.0 nM) was calculated by plotting the relative luminescence intensity of each group (n = 6) using GraphPad Prism 8.0 with nonlinear one-site binding regression. **g**) Normalized fluorescence intensity of hydrophobicity probe ANS after adding Aβs with or without CRANAD-17 (100 – 1600 nM). The slope is used as a hydrophobicity index, showing change of hydrophobicity upon mixing with CRANAD-17. **h**) Molecule docking of the binding poses of CRANAD-17 with Aβs (PDB: 5OQV). Protein surface is colored according to amino acid hydrophobicity (red: hydrophobic, blue: hydrophilic). The results suggest that CRANAD-17 binds to the hydrophobic moiety LVFF via the interaction of benzene and imidazole rings within the hydrophobic pocket of Aβs.

To further confirm whether CRANAD-17 could lead to changes of epitope hydrophobicity, fragment Aβ17-24 and full-length of Aβ were used in ANS (1-anilino-8-naphthalenesulfonate) fluorescence assay, which is commonly used for measuring protein surface hydrophobicity.^20, 21^ Normalized fluorescence intensity was plotted against protein concentration, and the slope was used as a hydrophobicity index to evaluate change. As shown in Fig.2g and Supplementary Fig.5, the hydrophobic index of Aβ significantly changed upon mixing with CRANAD-17, indicating alteration of epitope hydrophobicity. Similarly, the hydrophobic index of Aβ17-24 significantly changed in the presence of CRANAD-17 (Supplementary Fig.5). To further validate the hydrophobic interaction between Aβ and CRANAD-17, we performed computational simulation based on Aβ structural models 5OQV and 2LMO from the Protein Deposition Bank (PDB),^22^ and used ThT for comparison. Consistent with our previous NMR studies, both molecular docking data suggested that CRANAD-17 binds to the hydrophobic moiety LVFF via the interaction of benzene and imidazole rings within the hydrophobic pocket of Aβ (Fig.2h). In contrast, docking results of ThT showed dominant sites around ^10^YEVHHQK^16^ in 5OQV model and around C-terminal in 2LMO model (Supplementary Fig.6 and Supplementary Table 1). Taken together, CRANAD-17 could alter/edit the hydrophobicity of Aβ17-24 epitope to facilitate antibody binding.

The above results of CRANAD-17 were obtained from pure solution tests. However, it was not clear whether CRANAD-17 has similar editing effect on Aβs in a biologically relevant environment. In this regard, we used the media from 7PA2 cells, which is a familial APP mutation transfected cell line that secretes Aβ1-40 and Aβ1-42.^23^ Despite the low concentration of Aβ in cell media, the epitope editing effect could still be detected (Supplementary Fig. 7a,b). In addition, transgenic mice brain homogenate displayed enhanced readout after interacting with CRANAD-17 (Supplementary Fig. 7c,d). These data suggests that CRANAD-17 could execute its editing effect in biologically relevant environment.

Over the past several years, we have synthesized numerous curcumin analogues CRANAD-Xs.^24^ To investigate whether these analogues have similar a effect as CRANAD-17, we used the SPEED to test CRANAD-Xs (X= −3, −25, −44 and −102) (Supplementary Fig.8,9). We found that these compounds could alter the 4G8 antibody recognition and lead to increased chemilumiescence intensity. Interestingly, we found that CRANAD-102 was not able to edit the epitope from insoluble Aβ aggregates (Supplementary Fig.9e,f). This result is consistent with our previous report, in which we demonstrated that CRANAD-102 had good selectivity for soluble Aβs over insoluble Aβ aggregates.^24^

### Functional Studies for Epitope Editing

The charges on e-amino of lysines of Aβ, which are capable of forming salt bridges to stabilize mis-folded conformations, play an important role in Aβ aggregation and neurotoxicity.^9^ Previously, we found that 12-crown-4 showed decreased level of Aβ aggregates, confirmed by fluorescence assay and TEM observation.^13^ To further prove that the charge editing effect could break the salt bridge of Aβ and change protein aggregation behaviors, SDS-page gel studies were performed. As expected, the addition of 12-crown-4, 18-crown-6 and Orange G significantly changed aggregation behaviors of Aβs, as indicated by reduced aggregation of low molecular weight of Aβs (Supplementary Fig.10).

Previous studies indicated that Aβ species could initiate antibody production in vivo.^7, 25^ Importantly, microglia activation depends on Aβ conformation and core fragment/epitope of Aβ is related to neuroinflammation.^26^ We speculated that editing the hydrophobicity of epitope Aβ17-24 with CRANAD-17 could change immune-responses of Aβ and consequently alter pro-inflammatory properties of Aβ.

To this end, we determined the cytokine secretion of human microglia cells (HMC3) that were treated with Aβ in the presence or absence of CRANAD-17. As shown in Fig.3a-j, CRANAD-17 could significantly reduce the expression of various pro-inflammatory cytokines such as IL-4, −6 and −8, indicating excellent anti-inflammation capability of CRANAD-17 in HMC3 cells. Next, the level of expressed genes of HMC3 cells was determined by RNA-seq (Fig.3k and Supplementary Fig.11). The cells treated with Aβ oligomers showed higher genes expression related to type I interferon signaling pathway and IL-8 compared with control group. CRANAD-17 decreased the level of these pro-inflammatory related genes, further confirming the anti-inflammation capability of CRANAD-17 in HMC3 cells. In addition, we investigated the *in vivo* anti-inflammation effect of CRANAD-17 in 5xFAD transgenic mice, which were treated with CRANAD-17 for 3 weeks at the age of 9-month old, and sacrificed at the end of treatment. The brain slides were co-stained with anti-Aβ antibody 3D6 (for Aβ plaques) and microglia specific antibody IBA-1 (for microglia activation). We quantified the relative microglial activation using the intensity ratio of IBA-1 and 3D6 staining of the same plaques, and we found that CRANAD-17 treatment showed significantly decreased microglial activation (Fig.4a,b). In addition, we also found that the plaques were smaller after the treatment (Fig.4c). Taken together, our results suggested that epitope editing could lead to functional changes of target protein *in vitro* and *in vivo*.

**Fig.3.**
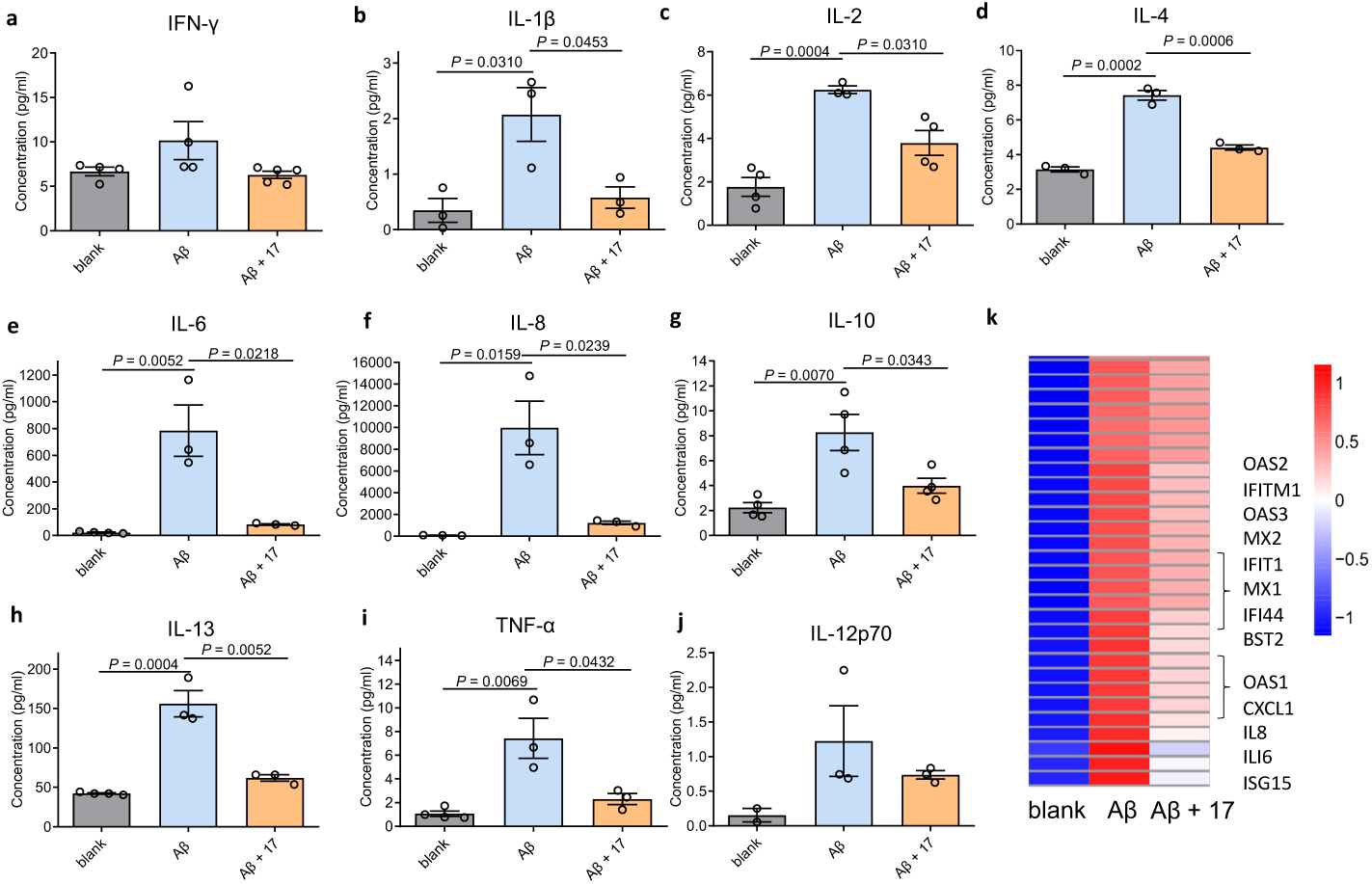
Anti-inflammation effect of CRANAD-17 in HMC3 cells. **a-j**) The changed levels of proinflammatory cytokines (pg/ml) secretion in human microglia cells (HMC3): cells (grey bar) were treated with Aβs with (orange bar) or without (blue bar) CRANAD-17 for 24 h and the cytokines in cell media were determined by meso scale discovery (MSD) assays (n ≥ 3). In the presence of CRANAD-17, the levels of most proinflammatory cytokines induced by Aβs, especially IL-6 (**e**) and IL-8 (**f**) were significantly decreased. Data are mean ± s.e.m. and *P* values were determined by two-tailed Student’s *t*-test. **k**) RNA-Seq analysis and representative hierarchical clustering heat map of differential expression genes from HMC3 cells with or without Aβs and CRANAD-17 treatment. The cells treated with Aβs showed highly expressed genes related with interferon signaling pathway and IL-8 compared to the control group. The level of these proinflammatory-related genes were apparently decreased with CRANAD-17 treatment. Color bar: large log10(FPKM+1) (red) and small log10(FPKM+1) (blue).

**Fig.4.**
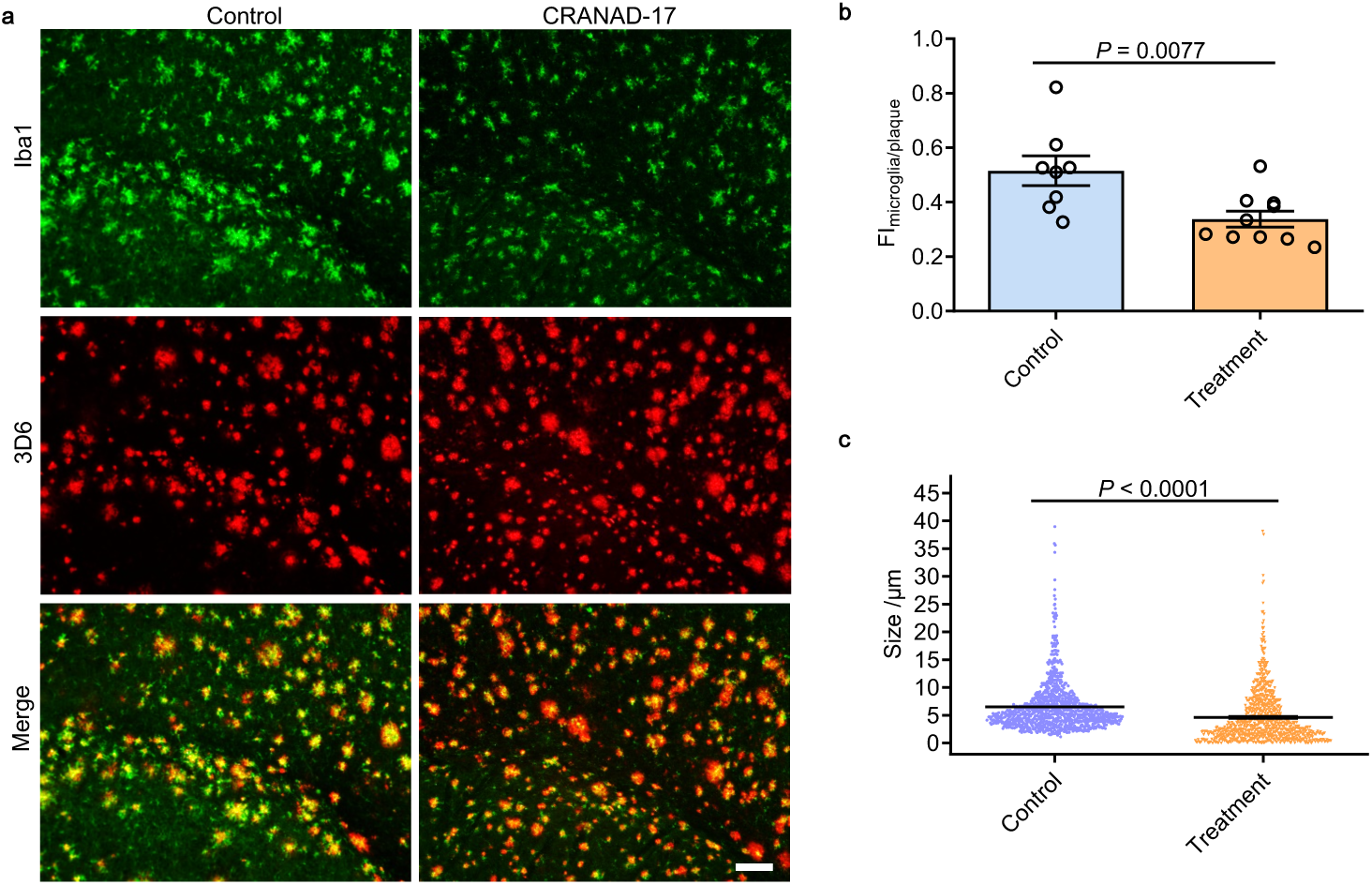
Anti-inflammation effect of CRANAD-17 in 5xFAD transgenic mice brains. **a**) representative immunofluorescence staining images of microglia cells (green) and Aβ plaques (red) stained by IBA-1 antibody and 3D6 antibody, respectively (scale bar = 50 μm). **b**) Quantitative analysis of the level of microglia activation. The average fluorescence intensity of microglia cells (n ≥ 250 in each tissue section) in mouse brain sections (n ≥ 8) were quantified and normalized with the fluorescence intensity of the corresponding background, showing significant decrease of microglia activation after CRANAD-17 treatment. **c**) The analysis of the size of Aβ plaques (n ≥ 2000), showing decreased size after CRANAD-17 treatment. Data in **b,c** are mean ± s.e.m. and *P* values were determined by two-tailed Student’s *t*-test.

### Discovery of Obatoclax as a Lead Imaging Ligand for Aβs via a Small Library Screening

To explore whether SPEED can be used to discover new leads for drug discovery and seeking imaging ligands for Aβs, we performed screening with a small library of 32 compounds that are FDA approved drugs or drug candidates. The full list of the compounds are shown in Supplemental Table 2 and Table 3. Indeed, the screening results showed that obatoclax, an inhibitor of Bcl-2 protein,^27^ and GNF-5837, a tropomyosin receptor kinase (Trk) inhibitor,^28^ showed significantly increased signal for Aβs with 4G8 antibody (Fig.5a,b). To confirm whether the obatoclax hit is true positive hit, we performed a concentration-dependent titration. As expected, obatoclax showed an excellent dose-dependence for Aβ binding (Fig.5c,d). To further validate that obatoclax is able to bind to the epitope of Aβ17-24, we performed molecular docking studies. The results confirmed that the interacting sites of obatoclax contain Aβ17-24. (Fig.5e, Supplementary Fig.6 and Supplementary Table 1).

**Fig.5.**
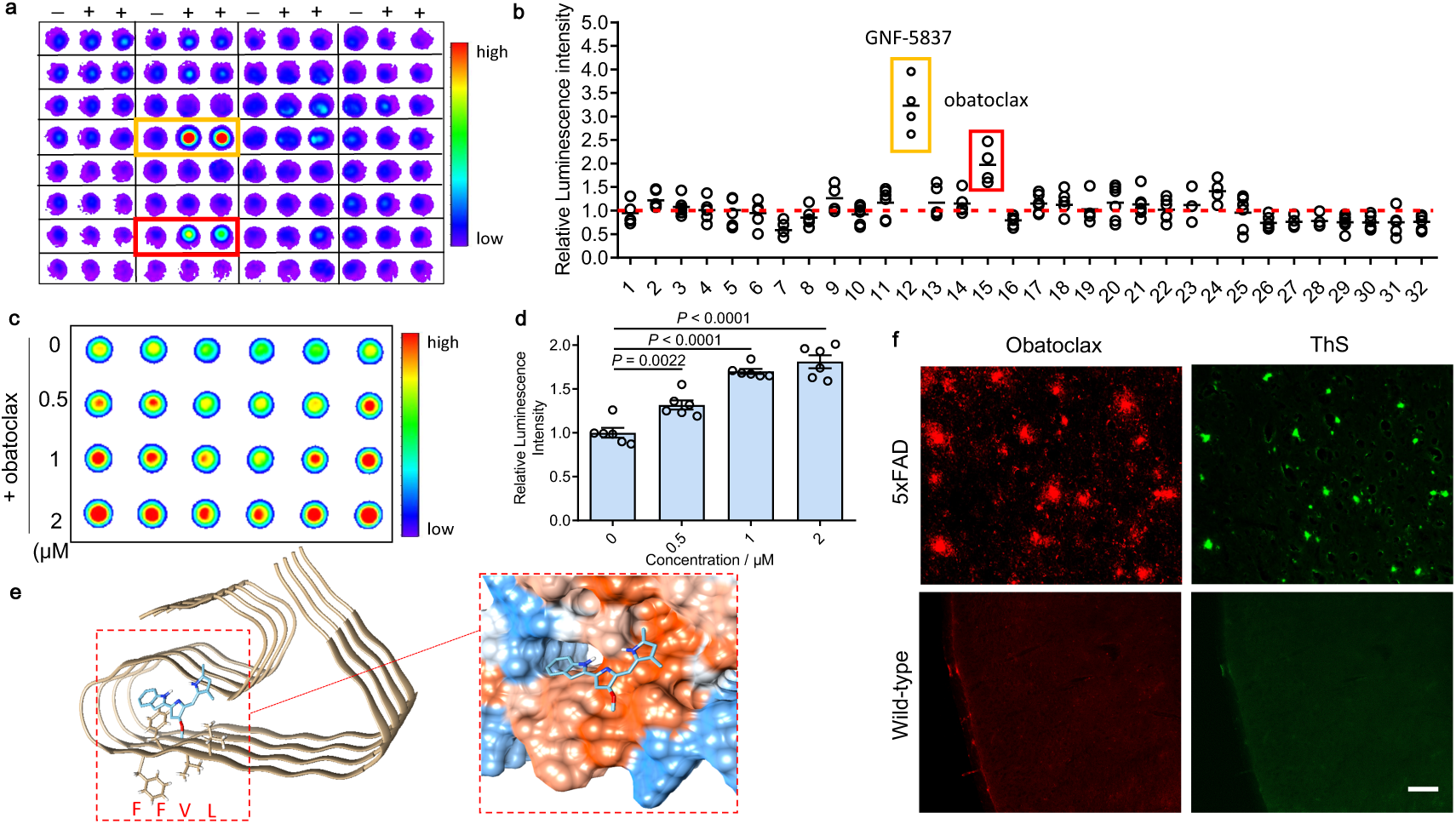
Small library SPEED test for seeking Aβ imaging agents. **a**) Representative SPEED screening results with a small library of 32 compounds (n = 2 in each assay, assay performed in duplicate). Obatoclax (red square) and GNF-5837 (orange square) showed obviously increased signal for Aβs with 4G8 antibody. **b**) Quantitative analysis of the screening results. Quantification was conducted using the ratio of the readout of each compound to that of blank control (red dashed line, defined as 1.0). Obatoclax and GNF-5837 were selected as hits (*P* < 0.0001 compared with blank control). **c,d**) SPEED assay (**c**) and quantitative analysis (**d**) of Aβs with different concentrations of obatoclax by using 4G8 antibody (n = 6). The results showed moderately increased signal for Aβs upon mixing with obatoclax, confirming the editing of epitope. **e**) Molecule docking of the binding poses of obatoclax with Aβs (PDB: 5OQV). Protein surface is colored according to amino acid hydrophobicity (red: hydrophobic, blue: hydrophilic). The interacting sites of obatoclax contain ^17^LVFFAEDV^24^ segment. **f**) Ex vivo histological staining of brain slices from a 9-month old 5xFAD mouse and wild-type mouse after 2mg/kg obatoclax administration. The brain sections of 5xFAD mouse treated with obatoclax showed substantial staining of Aβs that were confirmed by subsequent staining of 0.1% Thioflavin S, while wild-type mouse brain showed no labeling (scale bar = 50 μm). These data suggests obatoclax could potentially be used for fluorescence imaging of Aβs and the SPEED platform could be applied for seeking imaging probes.

Given the fluorescent properties of obatoclax, we then validated whether obatoclax could be used as a fluorescent imaging probe for Aβ species, we performed *ex vivo* histological observation of 5xFAD and wild-type mice brain. After intravenous injection of obatoclax, the mice brains were sliced and observed with fluorescence microscopy. The sections of 5xFAD mouse brain showed substantial staining of Aβ plaques that were further confirmed by subsequent ThS staining, while wild-type mouse brain showed no labeling (Fig.5f). These results indicated that obatoclax could potentially be used for fluorescence imaging Aβ plaques and the SPEED platform could be applied for seeking imaging probes.

We also noticed that GNF-5837 was orange and could be a potential fluorescent probe for Aβs. However, we found that it was not able to provide acceptable contrast for Aβ plaques in an AD mouse brain slice, which is likely due to the low quantum yield of GNF-5837. Therefore, no further action was taken for this compound.

### Application of SPEED for Other Targets

To further validate the applicability of the SPEED strategy, other targets were tested. Tau protein, which is a microtubule-associated protein with a critical role in several tauopathies include AD, Parkinson’s disease and Pick’s disease, was selected as a target protein.^29–31^ The VQIVYK segment of tau protein has been shown to drive tau aggregation and thus is considered an important target for seeking inhibitors for inhibiting tau fibrilization. However, the clear interaction mechanism between tau fibrils and inhibitor is still elusive. To test whether several commercially available tau inhibitors, tracers and dyes can alter the epitope VQIVYK segment, we selected ATPZ, T-807 and thioflavin S (ThS) to conduct the assay.^29–31^ As shown in Fig.6a,c, ThS showed significantly increased signal upon interacting with tau protein, which is in accordance with previous studies that indicate ThS could bind to the VQIVYK fragment.^31^ In constrast, the editing effect of ATPZ and T-807 on VQIVYK-contained epitope is negligible (Fig.6a-d), indicating these compounds may not have strong binding to this epitope.

**Fig.6.**
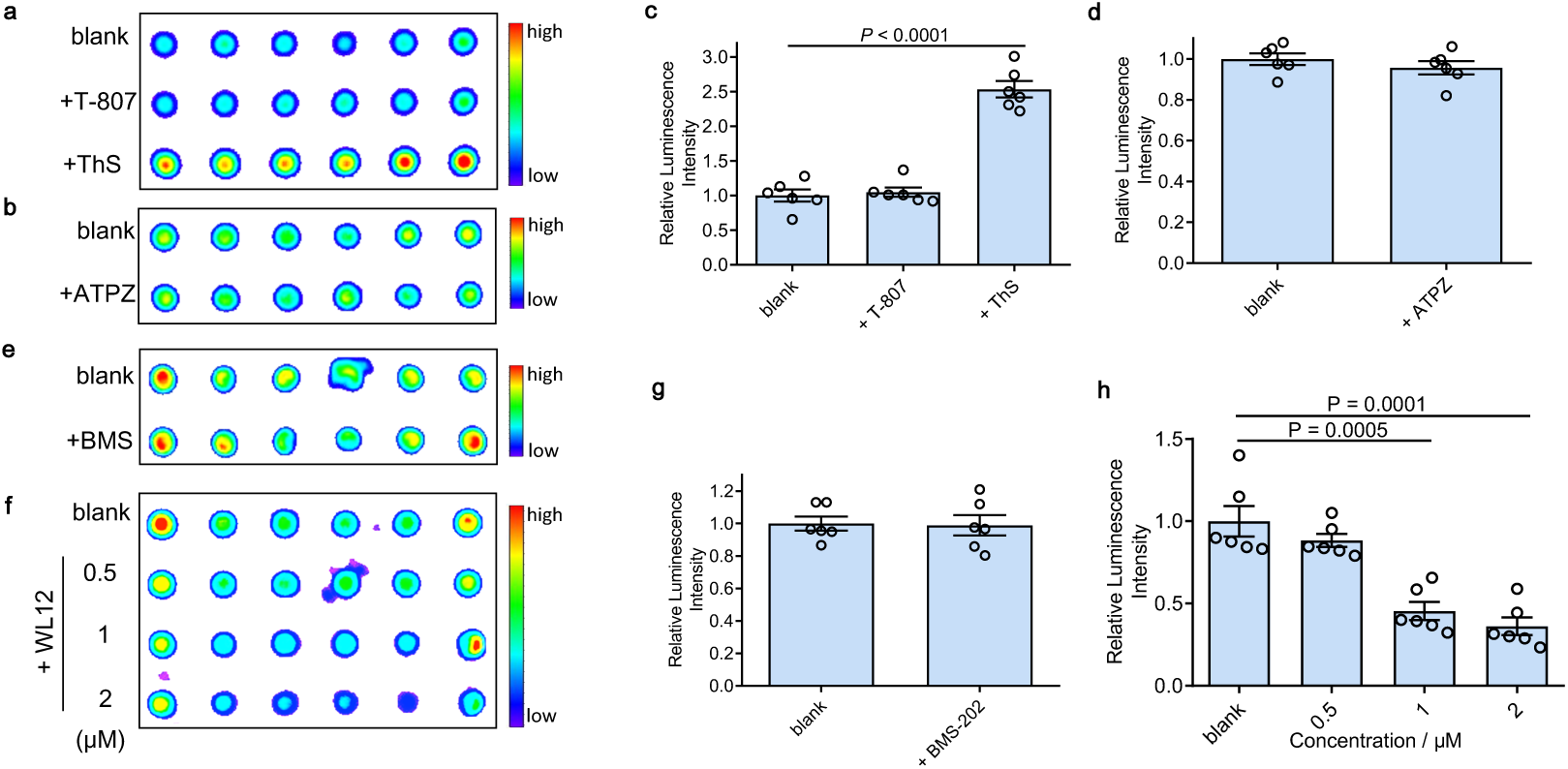
Application of SPEED for other targets. **a-d**) SPEED assays (**a,b**) and quantitative analysis (**c,d**) for full-length human tau protein treated with T-807, ATPZ and ThS using 8E6/C11 antibody, whose epitope lies within 209-224 a.a. of human tau (3-repeat isoform RD3) containing VQIVYK segment (n = 6). T-807 and ATPZ showed negligible signal change, suggesting the interacting sites of these compounds for tau protein are not within 209-224 a.a. segment. Only ThS showed significantly increased signal, which is in accordance with previous studies that indicate ThS could bind to the VQIVYK fragment. **e-h**) SPEED assays (**e,f**) and quantitative analysis (**g,h**) for PD-L1 protein with BMS-202 (**e,g**) and different concentrations of WL12 (**f,h**) using anti-PD-L1 antibody [28-8], whose epitope lies within 19-239 a.a. of PD-L1 (n = 6). The readout from WL12 treatment significantly decreased while BMS-202 displayed no obvious change compared to the control. Quantitative data are mean ± s.e.m. and *P* values were determined by two-tailed Student’s *t*-test.

To further expand the applicability of the SPEED platform, programmed death-ligand 1 (PD-L1) was selected as another model protein. PD-L1 is a membrane-bound protein expressed on the surface of tumor cells that inhibit T cell activation by binding to one of its complementary ligands PD-1. Recently, blocking the PD-1/PD-L1 pathway has become one of the most promising strategies in cancer therapy and several monoclonal antibodies are currently on the market or in the process of FDA approval.^32, 33^ Anti-PD-L1 antibody [28-8], which targets the specific extracellular domain of Phe19-Thr239 of PD-L1, and two classes of reported PD-L1 inhibitors include small molecule BMS-202 and peptide WL12 were selected for the testing.^32, 33^ As shown in Fig.6e-h, the readout of WL12 significantly decreased while BMS-202 displayed no obvious change compared to the blank control. This may arise from the flat and large interface of antibody-antigen interaction that makes it even more difficult for small molecular inhibitors to compete the binding.^34^ Several concentrations of WL12 were then used for the assay, further confirming the editing effect of epitope (Fig.6f,h). Taken together, our platform could be used to screen ligands for various targets and different classes of drug candidates, including both small molecules and peptide drugs.

## Discussion

Currently, only a few methods enable direct and label-free screening,^35, 36^ including drug affinity responsive target stability (DARTS),^37, 38^ and surface plasmon resonance (SPR) technology.^39^ In this report, we demonstrated that, as proof-of-concept, epitope editing could be used to seek ligands for a target of interest, which requires no modification/tagging and no extra equipment. In our exploratory investigation, several examples were successfully demonstrated, including Aβ peptides, tau and PD-L1 proteins. The molecular weights of these targets of interest span from 4KD to 55KD, which represents a wide range of proteinoid targets. We showed that this platform was not only suitable for targets related to Alzheimer’s disease, but also for cancer immunology-related protein PD-L1. Although our examples are related to proteinoids, we believe that our platform can be expanded into other non-proteinoid of targets, such as oligosaccharides, a very common category of antigens for immune reactions.

In the course of our studies, we discovered that CRANAD-Xs, obatoclax, GNF-5837 (Aβ) could enhance the readout signals. We speculated two possible reasons for this interesting phemonema: 1) as paratopes favor binding to aromatic rings from epitope via hydrophobic interactions,^6^ it was reasonable to speculate that these compounds could provide several aromatic rings within the epitope and thus enhance the binding to antibody, which is in accordance with recent discovery of protein-protein interaction (PPI) stabilizers that showed cooperative roles in PPI;^40, 41^ 2) our data suggests CRANAD-17 could change the hydrophobicity of epitope and it is possible that the binding may cause conformational change of the epitope and “open up” the epitope, which could be recognised by antibody as enhanced signal. Despite the elusive mechanism, the readout changes indeed indicate the altering/editing of epitope that could be used for drug screening.

Notably, this phenomenon also provided several important implications: 1) the binding of antigen to antibody could be enhanced via epitope editing, and the therapeutic effects of the antibody could be potentially boosted; 2) although CRANAD-X and Aβ antibody bind to the same epitope region, they are not in a competitive relationship, which differs from other drug screening methods based on inhibiting titration; and 3) cautious interpretation might be necessary for results from ELISA and dot blot if compound treatment was applied in the studies.

Our method based on the altering of epitopes opens a new avenue for seeking therapeutics and imaging lead ligands. This method possesses several advantages compared with previously reported methods: 1) the SPEED strategy does not require tagging of the antigens and/or drug candidates; 2) it can provide unambiguous information about the binding site of the hit, because of the specific recognition between antibody and epitope; 3) it could be feasible even for low affinity compounds as the readout relates to the degree of epitope change instead of binding affinity; 4) compatible for various model antigens include synthesized proteins, cell media and brain lysates; 5) simple utilization without pre-treatment that enables fast, convenient, cost efficient screening; and 6) quantifiable of binding affinity by titration assay. Here, we propose a few potential applications: 1) screening for site (epitope)-selective binding ligands. Most drug discovery starts with screening for pharmacological activity first, and the identification of binding site is subsequently accomplished by using other methods including computational tools and protein crystallization. However, for a wide spectrum of protein targets, certain sites within amino acid sequence have already been characterized as the major functional sites. For example, a PPI usually involves a few key residues called “hot spot” that contribute to the majority of the binding of protein-protein interface, and hence the hot spot is potentially important for drug discovery.^34^ Our platform could designate a specific site/epitope for screening related hits, thus could provide rationale-based screening to potentially decrease the laborious screening effort; 2) screening for molecules to manipulate immune responses. Immune responses could play double-edged roles, with a good role of fighting off invading organisms and a bad role of causing autoimmune diseases.^42^ Our platform could screen for molecules that may strengthen antibody-antigen interaction to avoid immune escape of antigens, or block epitopes that inhibit innate immune attack in autoimmune diseases. Also, this versatile platform could be used to screen epitope or antibody mimics; 3) screening for imaging agents. For the discovery of imaging agents, either for optical or radioactive imaging, competitive binding studies using a standard ligand are normally required. However, spectral overlap may occur during fluorescence competitive tests, and the use of radioactive ligands has considerable limitations. Moreover, it is possible that the standard ligand may have different binding sites from the tested compound, which could lead to false “negative” hits. In contrast, SPEED could serve as a more simple and versatile screening method for seeking imaging agents; 4) assisting in etiology elucidation. Our platform could also be utilized for functional site exploration, binding sites validation etc. 5) assisting in genome editing: although genome editing is a powerful tool to manipulate disease conditions, the gene associations of some diseases are still elusive, such as the majority of late onset Alzheimer’s disease;^43^ therefore, epitope editing (immune editing) could be an important complement in genome editing.^42^

The SPEED strategy also has certain limitations. For example, prior knowledge of epitope region within an antigen is needed and the corresponding antibodies should be available. However, as the utilization of antibody is rapidly developing in both clinical therapeutics and research applications, we believe this limitation could be overcome by advancements in antibody discovery and the epitope mapping process. Further efforts on SPEED would be highly desirable in characterizing the underlying mechanism, providing rational design criteria and directed screening for target proteins. Further research is currently underway in our group.

## Methods

### General Information

All reagents were commercial products and used without further purification. CRANAD-Xs compounds were synthesized according to our previously reported procedures and compounds with purities more than 95% determined by analytical HPLC, were used for further evaluation.^17, 24^ Synthetic Aβ1-40 peptide was purchased from rPeptide and different forms of Aβ1-40 including monomer, oligomer and aggregate were prepared by our previously reported methods.^17^ Acetylated-K16 Aβ1-40 was purchased from GenScript. Purified anti-β-Amyloid 1-16 (clone 6E10), 17-24 (clone 4G8) antibodies were purchased from BioLegend. Anti-Tau (3-repeat isoform RD3) antibody (clone 8E6/C11) was purchased from MilliporeSigma. Recombinant anti-PD-L1 antibody [28-8], recombinant human PD-L1 protein and recombinant human tau protein were purchased from Abcam. HRP-conjugated goat anti-Mouse IgG (H+L) secondary antibody was purchased from Invitrogen. 5xFAD and wild type mice were from Rudolph Tanzi Laboratory. All animal experiments were approved by the Institutional Animal Use and Care Committee (IACUC) at Massachusetts General Hospital, and carried out in accordance with the approved guidelines.

### Cell Culture

Human microglia (HMC3) cells and 7PA2 cells were cultured in Dulbecco’s Modified Eagle’s Medium/F-12 medium with high glucose (Gibco, USA) supplemented with 2 mM L-glutamine, 200 μg/ml G418, 100 IU/mL penicillin, and 100 μg/mL streptomycin at 37 °C in an atmosphere containing 5% CO2.

### A General SPEED Procedure with Purified Proteins

Nitrocellulose membrane (0.2 μm, Bio-Rad) was soaked in 0.01 M phosphate buffered saline (PBS) for 30 min before use. A solution of synthetic or recombinant protein (1 μM, final concentration) was mixed with drug/editor solution (2 μM in 10% DMSO/PBS, final concentration) or vehicle. After 1 h incubation at room temperature, 90 μL of mixture was immobilized on the membrane using a 96-well Bio-Dot Microfiltration Apparatus under vacuum. Each strips including 5-6 duplications were assigned to corresponding antibody recognition for epitope of interest (EOI). Briefly, the membrane was blocked with 5% nonfat milk for 1 h at room temperature and incubated with primary antibody (1:2000 diluted with 5% nonfat milk) at 4°C overnight. After washed with 0.1% Tween 20 in TBS (TBS-T) for 3 × 10 min, the membrane was incubated with HRP-conjugated secondary antibody (1:2000 diluted with 5% nonfat milk) for 1.5 h at room temperature and washed with TBS-T for 3 × 10 min. Visualization of EOI was performed with enhanced chemiluminescent reagent (Pierce ECL Western Blotting Substrate) using an IVIS®Spectrum imaging system (Perkin Elmer, Hopkinton, MA) with blocked excitation filter and opened emission filter. The chemiluminescence readout from control group (without tested drug/editor) was normalized to 1.0, and the editing effect for EOI was quantified using the ratio between two groups.

### SPEED with Cell Media and Brain Lysates

See supplementary information.

### Zeta Potential Test

Aβ1-40 aggregates (400 nM, final concentration) in 1 mL double distilled water were mixed with compound solution (800 nM, final concentration in distilled water) and incubated for 1 h at room temperature. The mixture was then pipetted into a folded capillary zeta cell (Malvern Panalytical) and the zeta potential was measured by using a Nano-sizer (Malvern, Nano-ZS).^13^

### ANS Fluorescence Assay

Aβ1-40 monomers (100-1600 nM, final concentration) were incubated with or without compound solution (100-1600 nM, final concentration) for 1 h and fluorescence intensity was measured by using a F-4500 fluorescence spectrometer (Hitachi, Japan) with excitation/emission = 350/525 nm). To the samples, 10 μL of 400 μM 1-anilinonaphthalene-8-sulfonic acid (ANS) was then added, incubated for 3 min, and subjected to the same fluorescence assay procedures. The final fluorescence intensity of samples were normalized by deducting the intensity of original samples without ANS treatment. The fluorescence intensity versus protein concentration (%) was plotted and slope (S0, a hydrophobic index) was calculated by linear regression analysis.^21^

### Docking Studies

The structures of the Ab1-40 fibrils were extracted from RCSB Protein Data Bank (PDB: 5OQV and 2LMO), followed by the refining of the molecular properties using Dock Prep function found in UCSF Chimera (UCSF Chimera, version 1.13.1) which included the addition of hydrogen atoms, deletion of solvent, and assigning of AMBER ff14SB force field for standard residues. The structure of compounds were generated on IQmol (IQmol, version 2.11), and minimized with UFF force field. The docking was performed using AutoDock Vina 1.1.2 visualized on UCSF Chimera with the default parameters. The receptor search volume was set to contain the entire Ab fibril based on their surface structure on UCSF Chimera. Each ligand docking returned the top 9 binding poses ordered based on score function and the best docking pose and scoring values was extracted. Each docking was repeated in triplcates to assure results. The hydrophobic surface was depicted by UCSF Chimera based on Kyte-Doolittle scale with blue being most hydrophilic, and red being most hydrophobic.^44^

### Proinflammatory Cytokines in HMC3 Cells

HMC3 cells were seeded at 4 × 10^5^ in a 6-well cell culture plate (Costar) for 24 h before being treated with Aβ1-42 oligomers (3 μM, final concentration) with or without compound solution (5 μM, final concentration in 1% DMSO). After 24 h incubation, the cell media was collected for MSD (meso scale discovery) assays with proinflammatory panel 1 (human) kit. By following the protocols from manufacturers, the concentration of ten major inflammatory cytokines from each sample was measured by the SECTOR® MSD instrument (USA) and calculated by MSD discovery workbench software.

### RNA-seq analysis of HMC3 Cells

The cells were seeded at 4 × 10^5^ in a 6-well cell culture plate (Costar) for 24 h before treated with Aβ1-42 oligomer (3 μM, final concentration) with or without compound solution (5 μM, final concentration in 1% DMSO). After 24 h incubation, the cell RNA were extracted and purified by using RNeasy Mini Kit (QIAGEN). The whole step of library construction, sequencing and bioinformatic analysis was performed at Novogene Bioinformatics Technology Co., Ltd. Briefly, the NEB library cDNA fragments of preferentially 150∼200 bp in length were constructed and purified with AMPure XP system (Beckman Coulter, Beverly, USA), followed by construction of strand specific library. The libraries were sequenced on HiSeq 2000 and HiSeq 2500 platform (Illumina, San Diego, USA). Differential expression analysis of two conditions/groups was performed using the DESeq2 R package. Hierarchical clustering analysis was carried out with the log10(FPKM+1) of union differential expression genes of all comparison groups under different experimental conditions.

### In Vivo Anti-inflammation Test in Transgenic Mice

9-month old 5xFAD transgenic mice were intravenously injected with CRANAD-17 solution (4 mg/kg; 15% DMSO 15% cremophor and 70% PBS) or vehicle twice a week. After being treated for 3 weeks, the brains of 5xFAD mice were removed and cut into 6-µm-thick slices using a CM 1950 cryostat (Leica, Germany). The brain tissues were subjected to immunostaining with anti-Iba1 antibody to visualize microglia and 3D6 to visualize Aβ by following routine immunostaining protocols. Images were captured by using a Nikon Eclipse 50i microscope. The fluorescence intensity and size of Aβ plaques and microglia cells were quantified by using ImageJ software.^45^

### SPEED for Small Library Screening

A small library of 32 compounds that are FDA approved drugs or drug candidates solutions (2 μM in 10% DMSO/PBS, final concentration) were mixed with synthetic Aβ protein solution (1 μM, final concentration). After 1 h incubation at room temperature, the mixture was subjected to routine SPEED test with LVFFAEDV as EOI (n = 2 in each assay, assay performed at least in duplicate, more information in Supplementary Table 2 and Table 3). The *P* values between control and tested group were determined by two-tailed Student’s *t*-test and the compounds with *P* < 0.0001 were selected as the hits.

### Ex Vivo Histological Staining

9-month old 5xFAD mice and wild-type mice were intravenously injected with 100 uL obatoclax solution (1mg/ml, 5% DMSO, 5% cremorpho1, 90% PBS) via tail vein and sacrificed at 60 minutes post-injection. The brains were harvested and cut into 10 μm slices and then co-stained with 0.1% Thioflavin S (50% ethanol). After washing the slices with 80% ethanol for 1min, 70% ethanol for 1min and distilled water for 2 × 1 min, the fluorescence images were captured by using a Nikon Eclipse 50i microscope.

### Statistical analyses

Quantitative data shown as mean ± s.e.m. were analyzed and presented with GraphPad Prism 8.0 software. *P* values were determined by unpaired two-tailed Student’s t-tests to evaluate the difference between two groups. The differences were considered significant when *P* ≤ 0.05.

## Supporting information

SPEED Supplemental Info

## Author contributions

C.R. designed the project. B.Z. developed and validated the screening platform. B.Z., C.Z, Y.L. and K.Y. contributed to cell studies. J.Y., R.T., X.S., S.H., C.Z., and K.R. contributed to animal studies. R.V. and Y.S. contributed to docking studies. W.Y. synthesized and characterized the compounds. B.Z. and C.W. contributed to the small molecule library screening. Y.L. contributed to protein purification. C.R. and B.Z. analyzed the results, prepared the manuscript, figures and supplementary information. C.R., C.Z, C.Y. and Z.Z. provided reagents, materials, and analysis tools. All authors contributed to discussion and editing of the manuscript.

