## Supplementary material for "A screening platform based on epitope editing for drug discovery": SPEED Supplemental Info

[1]Athinoula A. Martinos Center for Biomedical Imaging, Department of Radiology, Massachusetts General Hospital/Harvard Medical School, Room 2301, Building 149, Charlestown, Boston, Massachusetts 02129; [2]West China School of Pharmacy, Sichuan University, Chengdu, 610041, China; [3]Department of Chemistry and Biochemistry, University of Oklahoma, Norman, OK, 73019; [4]Genetics and Aging Research Unit, McCance Center for Brain Health, MassGeneral Institute for Neurodegenerative Disease, Department of Neurology, Massachusetts General Hospital/ Harvard Medical School, Charlestown, MA, USA; [5]Department of Systems Biology, Harvard Medical School, Boston, MA, United States.

**Experimental Section.**

**7PA2 Cell Culture.** The Chinese hamster ovary CHO cell line stably expressing *Indiana* mutation in APP, also known as 7PA2 cells, was cultured in Dulbecco's Modified Eagle's Medium/F-12 medium with high glucose (Gibco, USA) supplemented with 2 mM L-glutamine, 200 μg/ml G418, 100 IU/mL penicillin, and 100 μg/mL streptomycin at 37 °C in an atmosphere containing 5% $CO_2$.[1-3]

**Aβs concentration measurement in cells and mice brain.** 7PA2 cells were seeded at $4 \times 10^5$ in a 6-well cell culture plate (Costar). After 24 h incubation, the cell media was collected and Aβs ($A\beta_{1-40}$, $A\beta_{1-38}$ and $A\beta_{1-42}$) concentration were quantified using the MSD triplex assay via an electrochemiluminescence-based multi-array method.[4,5] The brain of APP/PS1 transgenic mouse was homogenated with RIPA lysis buffer and incubated on ice for 30 min, followed by centrifuging at 14000 rpm for 15 min at 4 °C. The Aβs ($A\beta_{1-40}$, $A\beta_{1-38}$ and $A\beta_{1-42}$) concentration in supernatants were analyzed with MSD triplex assay by following the manufacturer protocols.

**SPEED with Cell Media and Brain Lysates.** Nitrocellulose membrane (0.2 μm, Bio-Rad) was soaked in 0.01 M phosphate buffered saline (PBS) for 30 min before use. The cell media of 7PA2 cells and brain homogenate of APP/PS1 transgenic mouse with or without CRANAD-17 (1-50 nM in 10% DMSO/PBS, final concentration) were immobilized on the nitrocellulose membrane and subjected to routine SPEED test.

**Anti-aggregation Studies.** DMSO or compound solution (10 μM, final concentration) was added to a freshly prepared $A\beta_{1-40}$ or $A\beta_{1-42}$ solution in PBS (5 μM, final concentration). The resulting mixture was incubated at 37°C for 24 hours, and was then subjected to gel electrophoresis. Samples were separated on Novex™ 4-20% Tris-Glycine Mini Gels and stained using Pierce Silver Stain Kit (Thermo Fisher Scientific) according to the manufacturer's protocol. The images were acquired with IVIS®Spectrum and the greyscale intensity of each band (n = 3) was measured with Image J software.

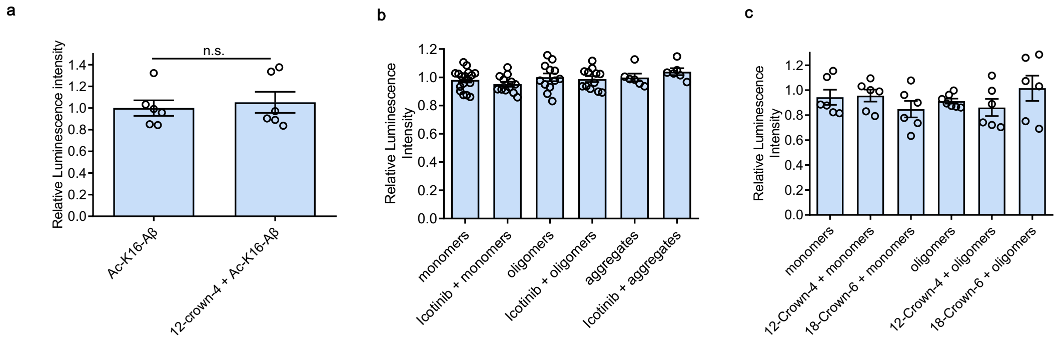

**Supplementary Figure 1 | Charge editing effect of crown ethers. a**). Relative luminescence intensity of SPEED assay for 1 μM acetylated-K16 Aβs (Ac-K16-Aβ) with or without 12-crown-4 (2 μM) by using 6E10 antibody (n = 6). The addition of 12-crown-4 showed negligible change of readout of Ac-K16-Aβ, further suggesting the altered charge on the site including K16. **b,c**) Relative luminescence intensity of SPEED assay for 1 μM Aβ monomers, oligomers and aggregates with or without 2 μM Icotinib (**b**), 12-crown-4 and 18-crown-6 (**c**) by using 6E10 antibody (n ≥ 6 for each group). Icotinib did not show significant editing effect. Crown ethers could only reduce the readout of aggregated Aβs but not monomeric and oligomeric Aβs.

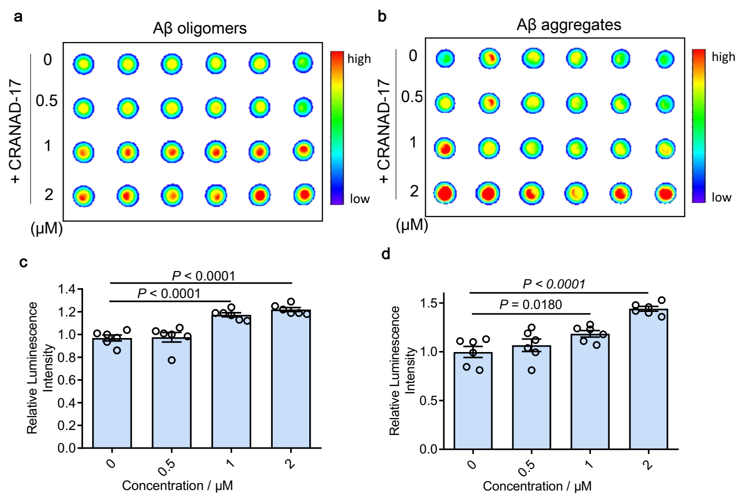

**Supplementary Figure 2 | Epitope editing effect of CRANAD-17 for different species of Aβs. a-d**) SPEED assays and quantitative data of 1 μM oligomers (**a,c**) and aggregates (**b,d**) with different concentrations of CRANAD-17 by using 4G8 antibody. The data showed significantly increased signal of all Aβ species after interacting with CRANAD-17 (The data of Aβ monomers were shown in Fig. 2**b,c**).

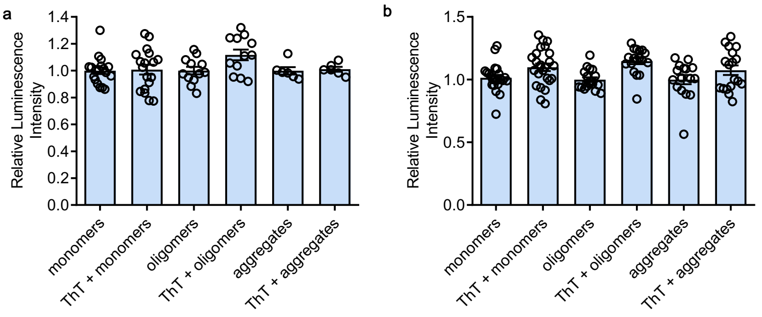

**Supplementary Figure 3 | Negligible epitope editing effect of ThT for Aβs detected by 4G8 and 6E10 antibodies.** Relative luminescence intensity of SPEED assays for 1 μM Aβ monomers, oligomers, aggregates with or without ThT (2 μM) by using 4G8 (**a**) and 6E10 (**b**) antibody (n ≥ 6 for each group). The data showed no apparent changes of Aβ species with and without ThT.

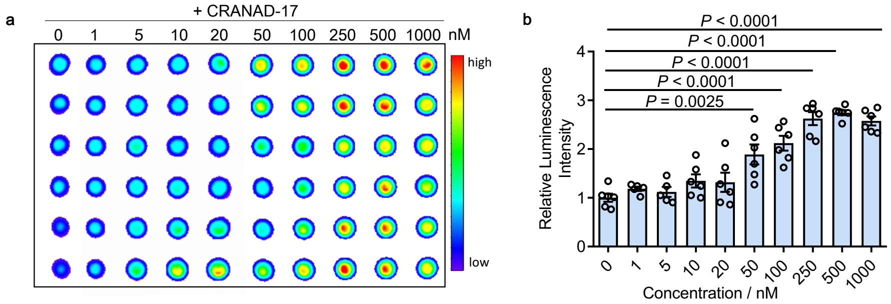

**Supplementary Figure 4 | Concentration dependent profile of CRANAD-17.** SPEED assay (**a**) and quantitative data (**b**) of Aβs upon mixing with CRANAD-17. The enhancement of readout was in a concentration-dependent manner.

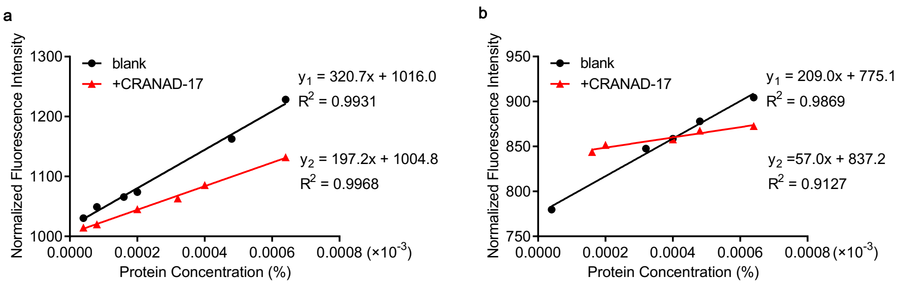

**Supplementary Figure 5 | Hydrophobicity change of Aβs upon interacting with CRANAD-17.** Normalized fluorescence intensity of hydrophobicity probe ANS after adding $A\beta_{1-40}$ (**a**) or $A\beta_{17-24}$ (**b**) monomers (100-1600 nM, final concentration) with or without CRANAD-17 (100 – 1600 nM). The slope is used as a hydrophobicity index, showing significant change upon mixing with CRANAD-17.

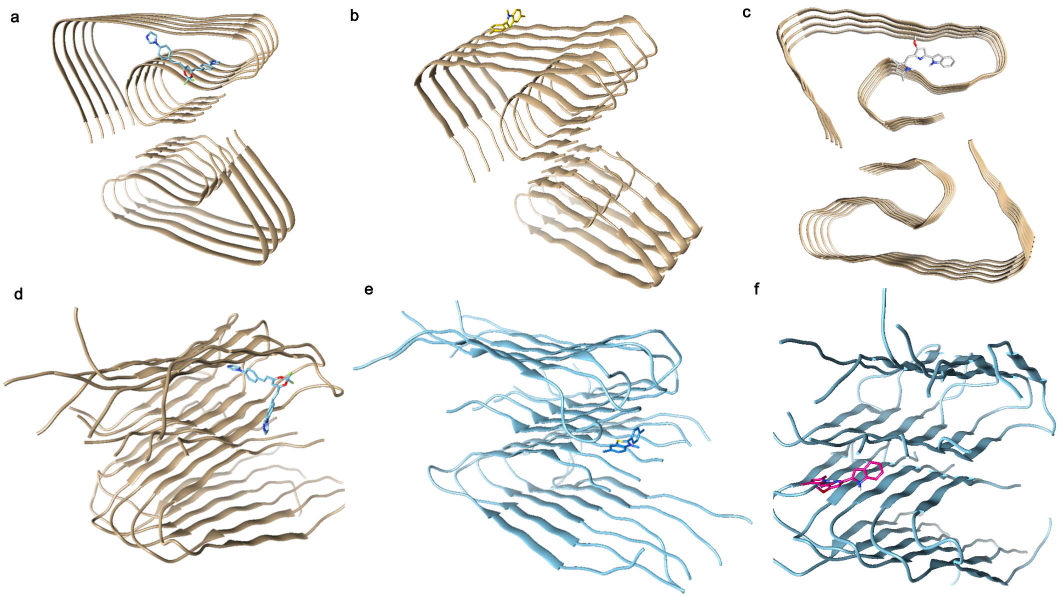

**Supplementary Figure 6 | Docking studies of CRANAD-17, Thioflavin T (ThT), and obatoclax.** Calculated binding poses of CRANAD-17 (**a,d**), ThT (**b,e**) and obatoclax (**c,f**) for $A\beta_{1\text{-}40}$ aggregates (PDB: 5OQV for **a-c** and PDB: 2LMO for **d-f**). CRANAD-17 displayed specific interaction within $^{17}$LVFFAEDV$^{24}$ segment in both models while ThT showed interaction at around $^{10}$YEVHHQK$^{16}$ for 5OQV model and around C-terminal in 2LMO model. Obatoclax showed specific interaction within $^{17}$LVFFAEDV$^{24}$ segment in 5OQV model and around C-terminal in 2LMO model.

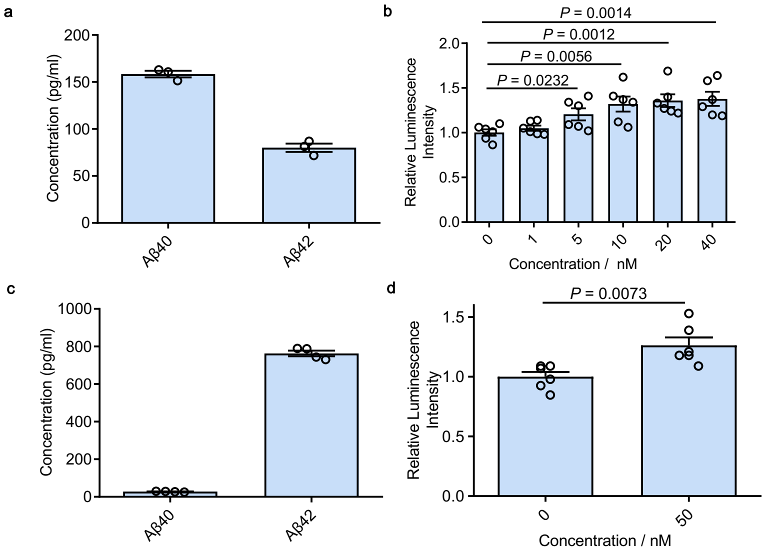

**Supplementary Figure 7 | Epitope editing effect of CRANAD-17 in biologically relevant environment. a,c**) $A\beta_{1-40}$ and $A\beta_{1-42}$ concentration of 7PA2 cell media (**a**) and 5xFAD transgenic mice brain homogenate (**c**) were measured by meso scale discovery test by using Aβ peptide panel 1 kit. **b,d**) Quantitative SPEED assay data of cell media (**b**) and transgenic mice brain homogenate (**d**) with or without CRANAD-17. The epitope editing effect could still be detected even at picogram concentrations of Aβs, indicating this method's high sensitivity.

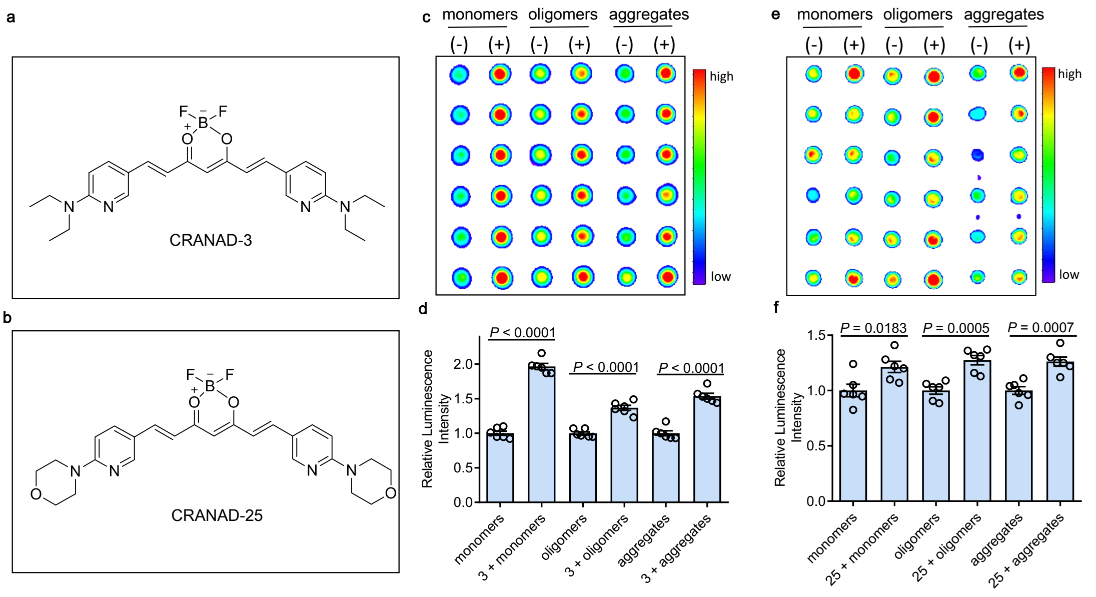

**Supplementary Figure 8 | Epitope editing effect of CRANAD-3 and -25 for Aβs.** **a,b**) Chemical structure of CRANAD-3 (**a**) and -25 (**b**). **c-f**) SPEED assays and quantitative data for 1 µM Aβ monomers, oligomers and aggregates with (+) or without (-) 2 µM of CRANAD-3 (**c,d**) and -25 (**e,f**) by using 4G8 antibody. CRANAD-3 and -25 could alter the 4G8 antibody recognition and lead to increased readout for all Aβ species.

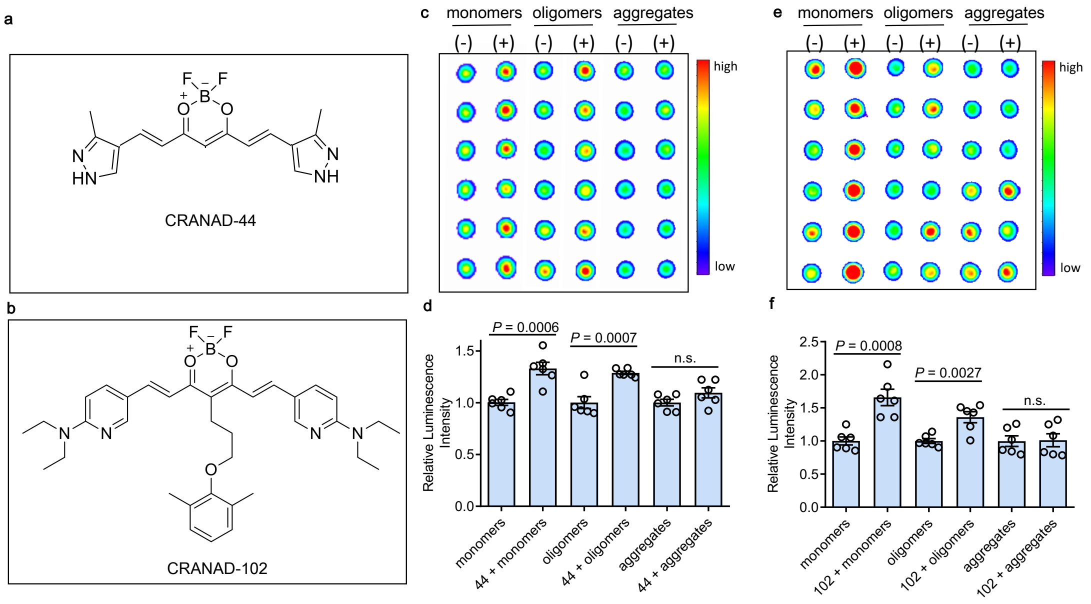

**Supplementary Figure 9 | Epitope editing effect of CRANAD-44 and -102 for Aβs.** **a,b**) Chemical structure of CRANAD-44 (**a**) and -102 (**b**). **c-f**) SPEED assays and quantitative data for 1 µM Aβ monomers, oligomers and aggregates with (+) or without (-) 2 µM of CRANAD-44 (**c,d**) and -102 (**e,f**) by using 4G8 antibody. CRANAD-44 showed increased readout for all Aβ species, while CRANAD-102 was not able to edit the epitope from insoluble Aβ aggregates. This result was consistent with our previous report, where we demonstrated that CRANAD-102 had good selectivity for soluble Aβs over insoluble Aβ aggregates.[6]

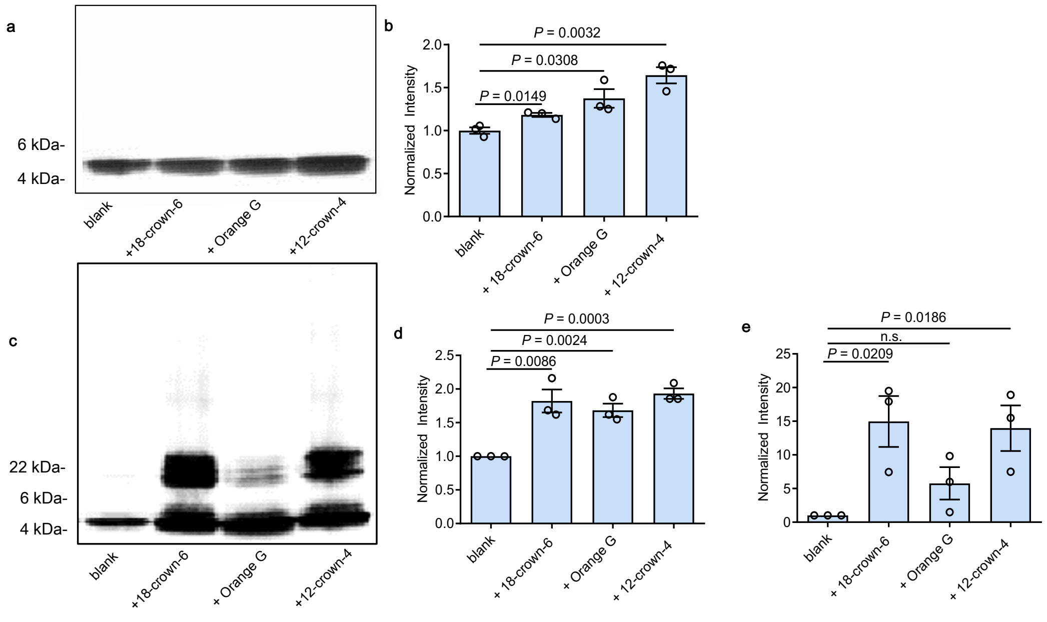

**Supplementary Figure 10 | Change of Aβ aggregation behaviors after charge editing.** Representative SDS-Page gel results of $A\beta_{1-40}$ (**a,b**) and $A\beta_{1-42}$ (**c-e**) with or without 18-crown-6, Orange G and 12-crown-4. Quantitative analysis (n = 3) of the greyscale intensity of monomers band (**b**,**d**) and low molecular weight oligomers band (**e**, only formed in $A\beta_{1-42}$ group) indicated decreased formation of insoluble Aβ aggregates, which are too large to enter the gel, after charge editing.

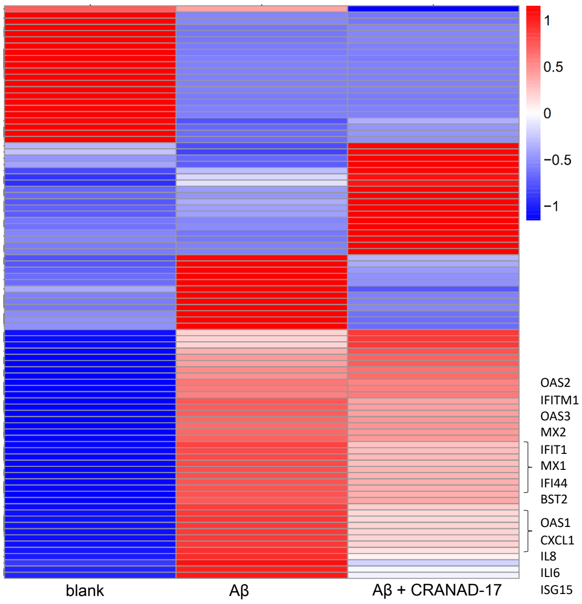

**Supplementary Figure 11 | Hierarchical clustering heat map of differential expression genes from human microglia cells (HMC3)**. HMC3 cells were treated with Aβ oligomers (3 μM) with or without CRANAD-17 (5 μM). The cells treated with Aβs showed higher gene expression related with type I interferon signaling pathway and IL-8 compared to the control group. The level of these proinflammatory-related genes decreased with CRANAD-17 treatment. Color bar: large log10(FPKM+1) (red) and small log10(FPKM+1) (blue).

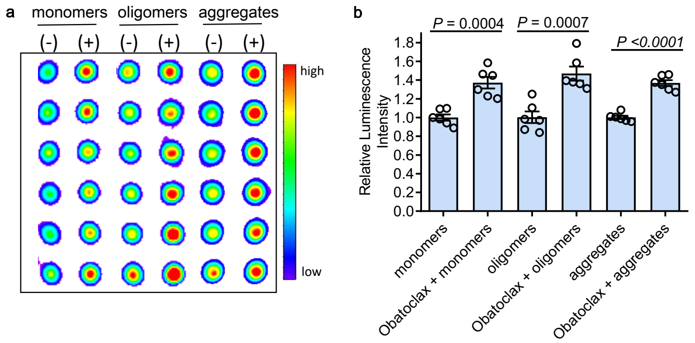

**Supplementary Figure 12 | Epitope editing effect of obatoclax for different species of Aβs.** SPEED assays (**a**) and quantitative data (**b**) of 1 μM Aβ monomers, oligomers and aggregates with (+) or without (-) 2 μM obatoclax by using 4G8 antibody. The data showed significantly increased signal from all Aβ species after interacting with obatoclax.

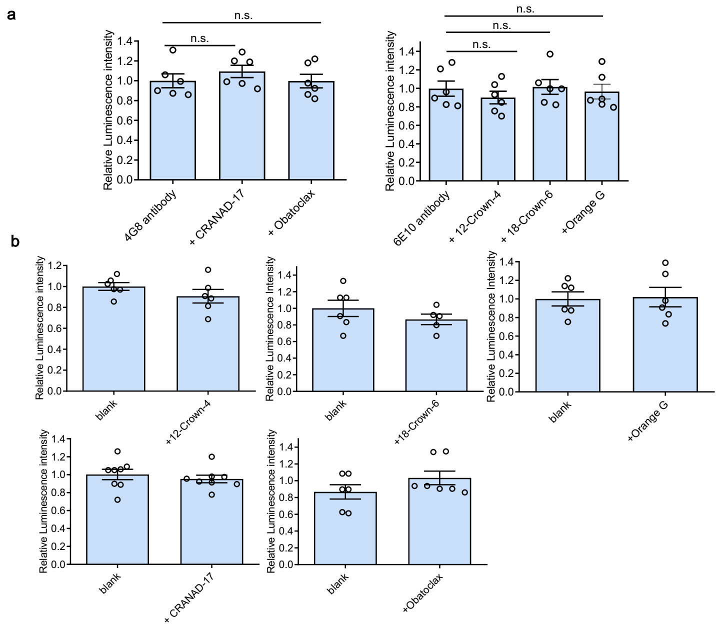

**Supplementary Figure 13 | No significant readout change of compounds with antibody only.** Relative readout of primary antibodies (**a**) and secondary antibodies (**b**) before and after adding the compounds used in the proof-of-concept studies. No significant change was found before and after adding the compounds, indicating the editing effects of compounds in SPEED assays are due to interaction with target proteins and not antibodies.

**Supplementary Table 1 | Autodock Vina Score of Compounds upon Interacting with Aβs.**

| PDB code | | CRANAD-17 | ThT | Obatoclax |
|---|---|---|---|---|
| 5OQV | Scores[a] | -9.4 | -7.2 | -7.3 |
| | a.a. segment[b] | [17]LVFFAEDV GSNKGAI[31] | [10]YEVHHQ KLV[18] | [17]LVFFAED VGSNKGAIIG[33] |
| 2LMO | Scores[a] | -10.1 | -7.9 | -8.6 |
| | a.a. segment[b] | [17]LVFFAEDV GSNKGAIIGLM[35] | [36]VGGVV[40] & [25]GSNKGAII[32] | [36]VGGVV[40] & [25]GSNKGAII[32] |

[a]Docking scores (kcal/mol) were calculated by Autodock vina. [b]Calculated interacting sites within Aβ model.

**Supplementary Table 2 | 96-well Plate Layout of Small Compound Library Based SPEED Screening**

| | 1 | 2 | 3 | 4 | 5 | 6 | 7 | 8 | 9 | 10 | 11 | 12 |
|---|---|---|---|---|---|---|---|---|---|---|---|---|
| A | blank | 1 | 1 | blank | 9 | 9 | blank | 17 | 17 | blank | 25 | 25 |
| B | blank | 2 | 2 | blank | 10 | 10 | blank | 18 | 18 | blank | 26 | 26 |
| C | blank | 3 | 3 | blank | 11 | 11 | blank | 19 | 19 | blank | 27 | 27 |
| D | blank | 4 | 4 | blank | 12 | 12 | blank | 20 | 20 | blank | 28 | 28 |
| E | blank | 5 | 5 | blank | 13 | 13 | blank | 21 | 21 | blank | 29 | 29 |
| F | blank | 6 | 6 | blank | 14 | 14 | blank | 22 | 22 | blank | 30 | 30 |
| G | blank | 7 | 7 | blank | 15 | 15 | blank | 23 | 23 | blank | 31 | 31 |
| H | blank | 8 | 8 | blank | 16 | 16 | blank | 24 | 24 | blank | 32 | 32 |

**Supplementary Table 3 | Sample Information of Small Compound Library Based SPEED Screening**

| | | | |
|---|---|---|---|
| 1 | 2 | 3 | 4 |
| Tetracycline | LG 100754 | BMS-202 | Phloretin |
| 5 | 6 | 7 | 8 |
| UVI 3003 | EF-24 | Chidamide | Panobinostat |
| 9 | 10 | 11 | 12 |
| JZL 195 | Martinostat | Lacmoid | GNF 5837 |
| 13 | 14 | 15 | 16 |
| GW 441756 | Rosiglitazone | Obatoclax | CN- Selisistat |
| 17 | 18 | 19 | 20 |
| F- Selisistat | IBET726 | CF53 | Selisistat |
| 21 | 22 | 23 | 24 |
| SR 17018 | YNT-185 | JQ-1 | S 02 |
| 25 | 26 | 27 | 28 |
| AX 024 | Mivebresib | Cutamesine | Necrostatin-1 |
| 29 | 30 | 31 | 32 |
| PD 144418 | EI 1 | C- Selisistat | SAHA |
